# Linking influenza virus evolution within and between human hosts

**DOI:** 10.1101/812016

**Authors:** Katherine S. Xue, Jesse D. Bloom

## Abstract

Influenza viruses rapidly diversify within individual human infections. Several recent studies have deep-sequenced clinical influenza infections to identify viral variation within hosts, but it remains unclear how within-host mutations fare in the global viral population. Here, we compare viral variation within and between hosts to link influenza’s evolutionary dynamics across scales. Synonymous sites evolve at similar rates at both scales, indicating that global evolution at these putatively neutral sites results from the accumulation of within-host variation. However, nonsynonymous mutations are depleted in global viral populations compared to within hosts, suggesting that selection purges many of the protein-altering changes that arise within hosts. The exception is at antigenic sites, where selection detectably favors nonsynonymous mutations at the global scale, but not within hosts. These results suggest that selection against deleterious mutations and selection for antigenic change are the main forces that transform influenza’s within-host genetic variation into global evolution.

## Introduction

As influenza viruses replicate within infected hosts, they quickly mutate into genetically diverse populations. A small proportion of within-host variants transmit between individuals (McCrone and Lauring, 2018; McCrone et al., 2018), and some transmitted variants continue to spread from person to person to circulate globally, and even reach fixation (Alizon et al., 2011; Mideo et al., 2008; Xue et al., 2018)(Figure 1). Influenza virus’s genetic variation within hosts therefore provides the material for its rapid global evolution.

**Figure 1.**
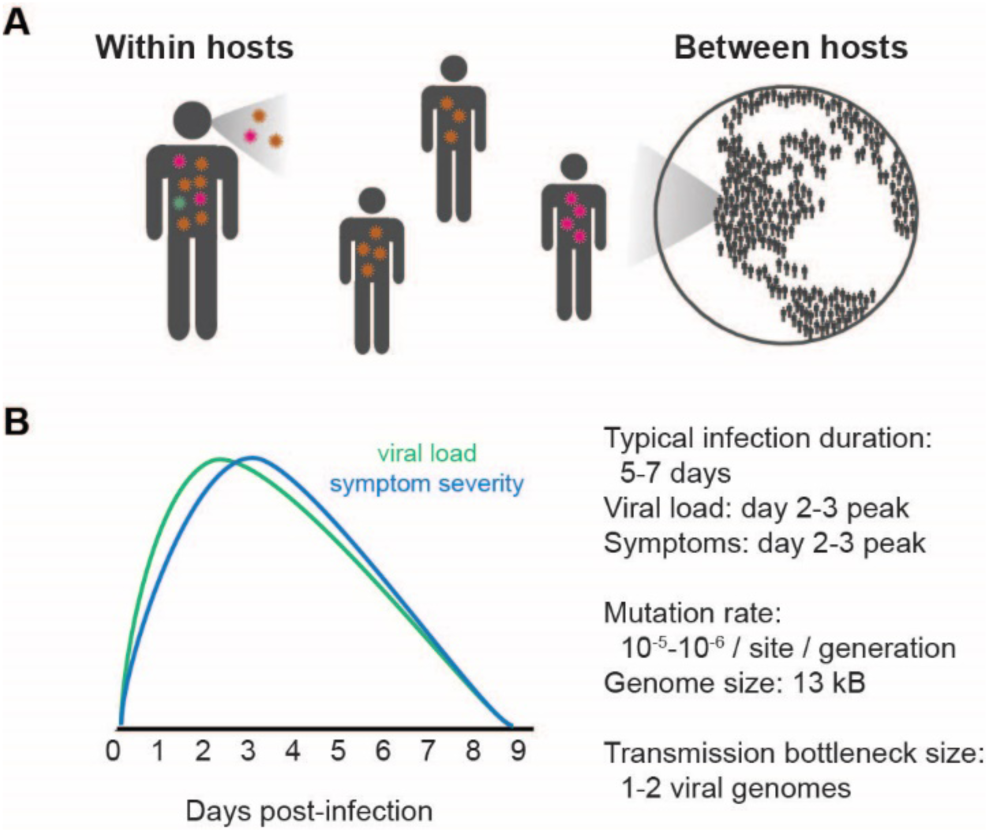
Influenza virus evolves at within- and between-host (or global) evolutionary scales. **A)** Viral mutations that arise within hosts can transmit from person to person and eventually contribute to global viral evolution. **B)** Key parameters affecting influenza’s evolution within hosts. Estimates of infection duration, viral load, and symptom severity are summarized from meta-analyses of volunteer challenge studies (Carrat et al., 2008) and mathematical models of viral infection (Baccam et al., 2006; Beauchemin and Handel, 2011). Viral mutation rates are summarized from studies by (Bloom, 2014; Nobusawa and Sato, 2006; Sanjuán et al., 2010; Suarez-Lopez and Ortin, 1994). Transmission bottleneck size has recently been estimated from deep sequencing of viral samples from household transmission pairs (McCrone et al., 2018).

Several recent studies have used deep sequencing to identify within-host mutations in hundreds of clinical influenza infections (Debbink et al., 2017; Dinis et al., 2016; McCrone et al., 2018). However, it remains unclear what role these within-host mutations play in the global evolution of influenza virus. The same within-host mutations are only rarely observed in different individuals, and mutations that reach detectable frequencies at the global scale are not notably more common than other mutations within hosts (Debbink et al., 2017; McCrone et al., 2018). Specifically, although influenza virus displays rapid antigenic evolution on the global scale, antigenic variants are present at low frequencies within hosts (Dinis et al., 2016; McCrone et al., 2018). New quantitative approaches are therefore needed to understand how within-host viral variation is transformed into global evolution.

Here, we compare the genetic variation of H3N2 influenza virus within and between human hosts to infer the fates of within-host mutations in the global viral population. Individual within-host mutations are challenging to track on a global scale, but by comparing large numbers of genetic variants identified within and between hosts, we can infer the evolutionary forces that act on different classes of mutations. We analyze within-host viral variation in 308 acute influenza infections from three deep-sequencing datasets (Debbink et al., 2017; Dinis et al., 2016; McCrone et al., 2018), and we calculate rates of evolution within and between hosts to determine how selection and genetic drift shape viral evolution. Synonymous mutations accumulate at similar rates within and between hosts, but nonsynonymous mutations are less prevalent globally than they are within hosts across most of the influenza genome. These observations suggest that many nonsynonymous mutations that reach detectable frequencies within hosts are later purged from the global influenza population. In antigenic sites, however, nonsynonymous mutations accumulate more rapidly on a global scale than they do within hosts, suggesting that antigenic selection primarily takes place between hosts. Our results show that influenza populations within hosts are dominated by transient, deleterious mutations which are later eliminated at transmission and the early stages of global evolution. Selection against these deleterious mutations and selection for antigenic change are the main forces that transform influenza’s within-host variation into global evolution.

## Results

### Rates of influenza virus evolution within hosts

Evolutionary rates provide a simple, quantitative framework for comparing viral variation across scales. Under neutral evolution, we expect within-host and global evolutionary rates to be identical, and deviations from these neutral expectations shed light on how selection acts across evolutionary scales. For instance, HIV and hepatitis C virus (HCV) both evolve more rapidly within than between hosts, probably because viruses acquire adaptations to specific hosts that often revert after transmission (Alizon and Fraser, 2013; Gray et al., 2011; Herbeck et al., 2006; Lemey et al., 2006; Lythgoe and Fraser, 2012; Raghwani et al., 2018; Zanini et al., 2015).

We sought to calculate the within-host and global evolutionary rates of influenza virus. Influenza’s global evolutionary rate is easily estimated from phylogenies of patient consensus sequences, which represent transmitted strains (Rambaut et al., 2008), but evolutionary rates during acute infections are more challenging to calculate. Several studies have estimated within-host evolutionary rates for chronic viruses like HIV and HCV by sequencing longitudinal viral samples (Alizon and Fraser, 2013; Gray et al., 2011; Lemey et al., 2006; Raghwani et al., 2018). However, longitudinal samples are difficult to collect for viruses like influenza that cause acute infections, and acute infections also provide limited time for genetic diversity to accumulate. Errors that arise in library preparation and sequencing are often present at similar frequencies to genuine within-host genetic variation in acute viral infections, making it challenging to accurately estimate within-host evolutionary parameters (Illingworth et al., 2017; McCrone and Lauring, 2016; Zhao and Illingworth, 2019).

To overcome these challenges, we developed a method to estimate rates of within-host evolution from deep sequencing of acute viral infections. In brief, we identified within-host mutations that were present in at least 0.5% of sequencing reads (Figure 2**, Materials and methods**). We calculated the total genetic divergence in each viral sample by summing the frequencies of within-host mutations. We then normalized divergence to the number of available sites in the genome and time since the infection started (Figure S1) to estimate a rate of evolution per site, per day (Figure 3).

**Figure 2.**
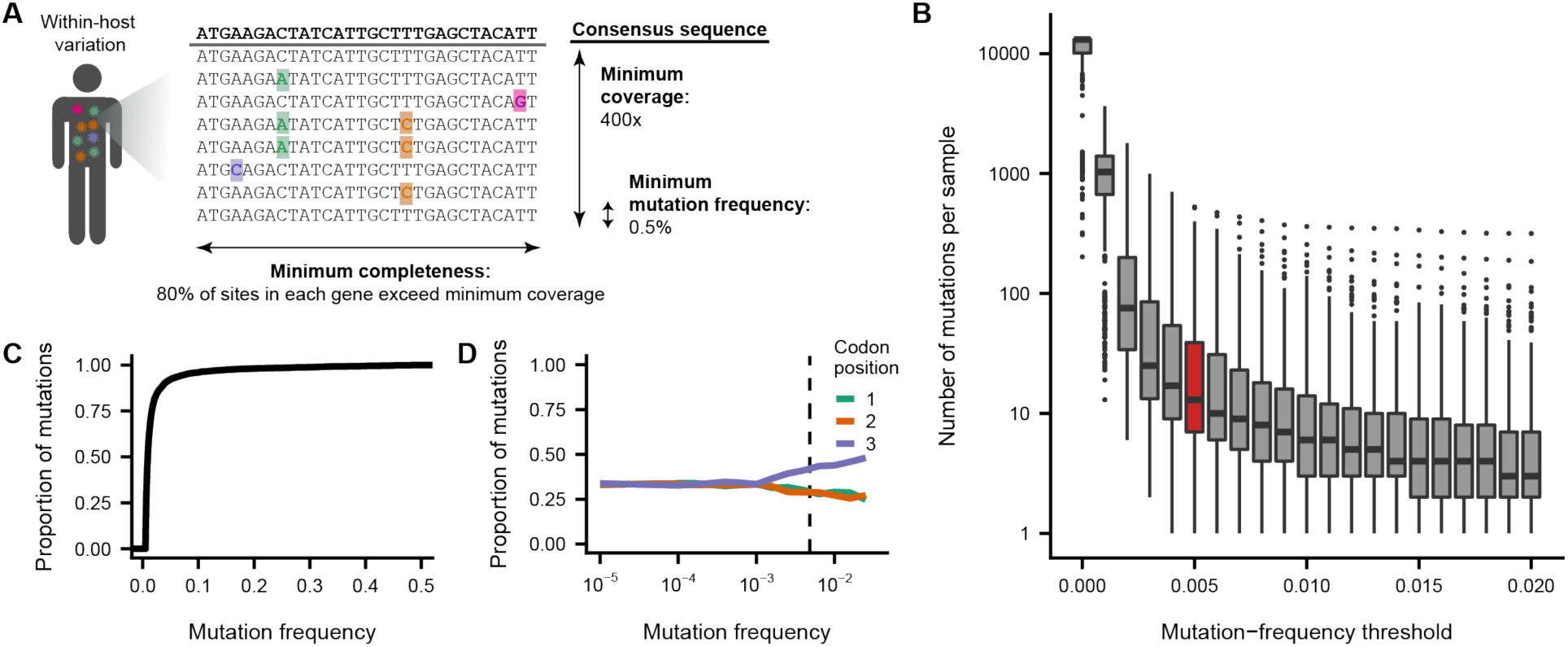
Within-host mutation calling criteria. **A)** Within-host mutations were called when a non-consensus base reached a frequency of 0.5% at a site with at least 400x sequencing coverage. We analyzed only viral samples in which >80% of the sites in each sequenced gene had ≥400x coverage. **B)** The number of mutations called in each viral sample declines as the minimum mutation-frequency threshold increases. Outlier samples with high within-host variation were included in this plot but excluded from subsequent analyses (Figure S3). **C)** Most within-host mutations called above a frequency of 0.5% are present at <5% frequency. **D)** Within-host mutations at higher frequencies are more likely to be located at the third codon position. Sequencing errors, which occur at low frequencies, are expected to be evenly distributed across all three codon positions. In contrast, high-frequency mutations are disproportionately located at the third codon position, where mutations are often nonsynonymous, suggesting that mutations at higher frequencies are more likely to represent true viral mutations that have experienced purifying selection. The dashed line indicates the 0.5% mutation-frequency threshold used in this study.

**Figure 3.**
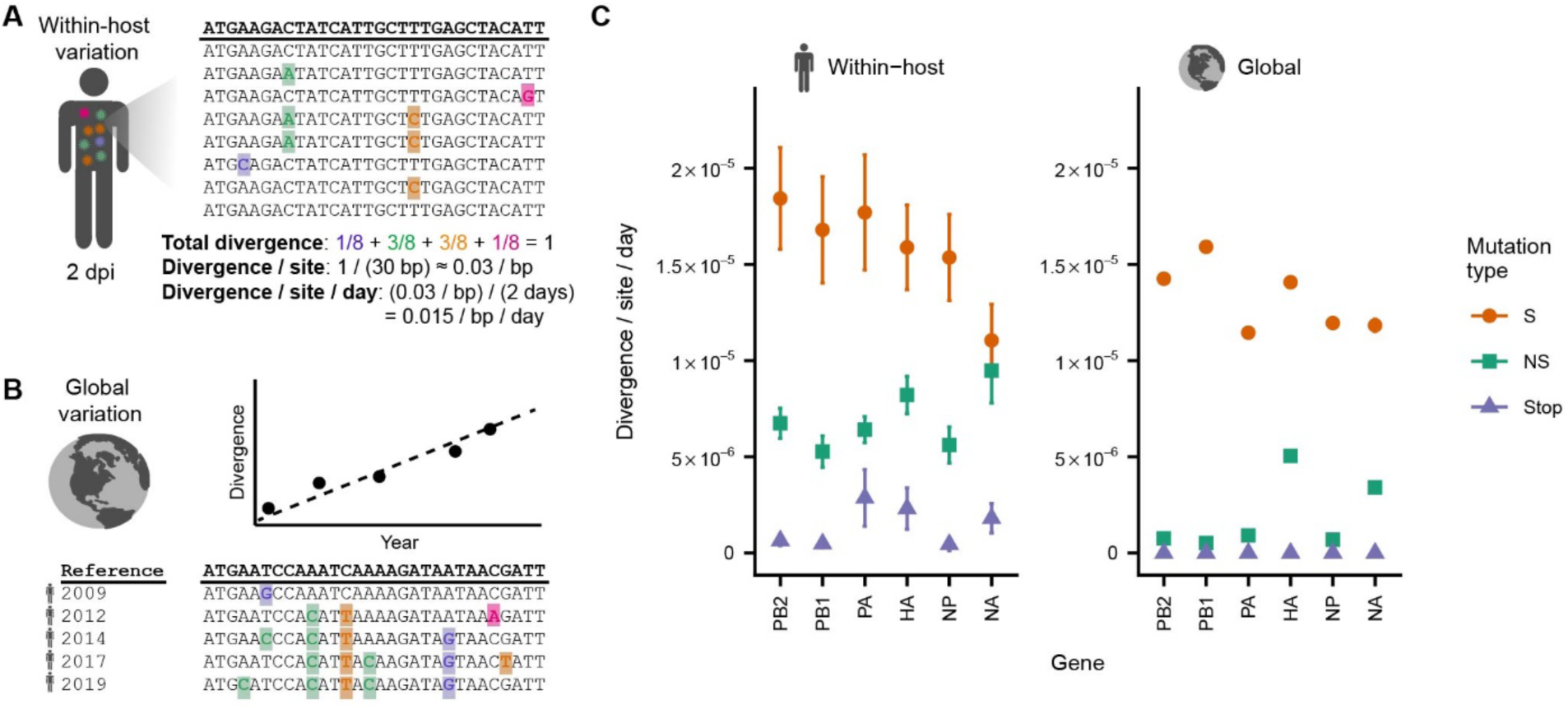
Within-host and global evolutionary rates reveal selective pressures acting on influenza virus across evolutionary scales. **A)** Within-host evolutionary rates were calculated by normalizing the total divergence of each within-host viral population to the number of sites and the time elapsed since the infection began. **B)** Global evolutionary rates were calculated using a molecular-clock method by performing a linear regression of divergence per site versus time. **C)** Within-host and global evolutionary rates of H3N2 influenza virus at synonymous (S), nonsynonymous (NS), and stop-codon (Stop) sites. Rates of synonymous evolution are broadly similar at the within-host and global scale, but nonsynonymous mutations accumulate more rapidly within hosts than globally. Within-host rates are shown as the mean and standard error of all patient infections sequenced in the three datasets analyzed after removing outlier samples (see **Materials and methods**) (Debbink et al., 2017; Dinis et al., 2016; McCrone et al., 2018). Global rates are shown as the mean and standard error calculated through linear regression (the standard errors are smaller than the point sizes). Global rates of stop-codon evolution are zero because no stop codons are observed in patient consensus sequences.

We tested the sensitivity of this method to common technical considerations. Estimates of evolutionary rates can be influenced by the frequency threshold used to identify within-host variation (Gallet et al., 2017; Grubaugh et al., 2019; McCrone and Lauring, 2016). We focused on within-host variants present in at least 0.5% of sequencing reads at a site (Figure 2A). At lower mutation-frequency thresholds, more mutations are identified in each viral sample (Figure 2B), but many of these putative mutations result from errors in library preparation and sequencing. Higher mutation-frequency thresholds limit the influence of false-positive mutations but can also exclude true within-host variation, most of which is present at low frequencies (Figure 2C). Our preferred mutation-frequency threshold of 0.5% is relatively permissive but comfortably exceeds the 0.1% threshold above which true variants begin to exceed sequencing errors, based on the proportion of putative mutations present at each codon position (Figure 2D)(Dyrdak et al., 2019). Nevertheless, we also calculated within-host evolutionary rates at different mutation-frequency thresholds and found that the relative evolutionary rates of different genes and site classes remained consistent (Figure S2).

In calculating within-host evolutionary rates, we sought to limit the influence of outlier samples with unusually high within-host variation (Figure 2B, Figure S3). This high variation can arise for biological reasons like co-infection with two distinct viral strains, or for technical reasons like poor sample quality or low viral load (McCrone and Lauring, 2016; McCrone et al., 2018). However, both sources of variation artificially inflate estimates of how quickly viruses evolve within patient infections. To minimize the influence of outlier samples, we ranked all samples based on the number of within-host mutations and excluded the top 10% of samples from each study. We also excluded samples that did not have at least 400x sequencing coverage in at least 80% of the sites in each gene.

We calculated rates of within-host influenza evolution from deep-sequencing data in three published studies that together represented 308 acute H3N2 influenza infections (Figure 3, Figure S3)(Debbink et al., 2017; Dinis et al., 2016; McCrone et al., 2018). We estimated evolutionary rates separately for synonymous, nonsynonymous, and stop-codon (nonsense) mutations in each patient and each viral gene, and we averaged the rates estimated for each patient to calculate a single rate of evolution for each gene and mutation type. Nonsynonymous mutations are more common than synonymous mutations due to the structure of the genetic code, so we normalized evolutionary rates to the number of possible sites for each mutation type (**Materials and methods**). We limited our analysis to the six longest of the eight influenza virus genes for two reasons. First, there is less sampling noise in the estimates of evolutionary rates for longer genes because there are more sites. Second, the two shortest influenza genes have alternatively spliced and partially overlapping reading frames that complicate annotation of mutations as nonsynonymous or synonymous.

To assess sources of variation in our estimates, we calculated evolutionary rates separately for each deep-sequencing dataset (Figure S4). We also used a second method to estimate evolutionary rates, performing linear regression of viral divergence by the time since the infection started (Figure S5). In both cases, the resulting evolutionary rates supported the qualitative conclusions described below.

**Figure 4.**
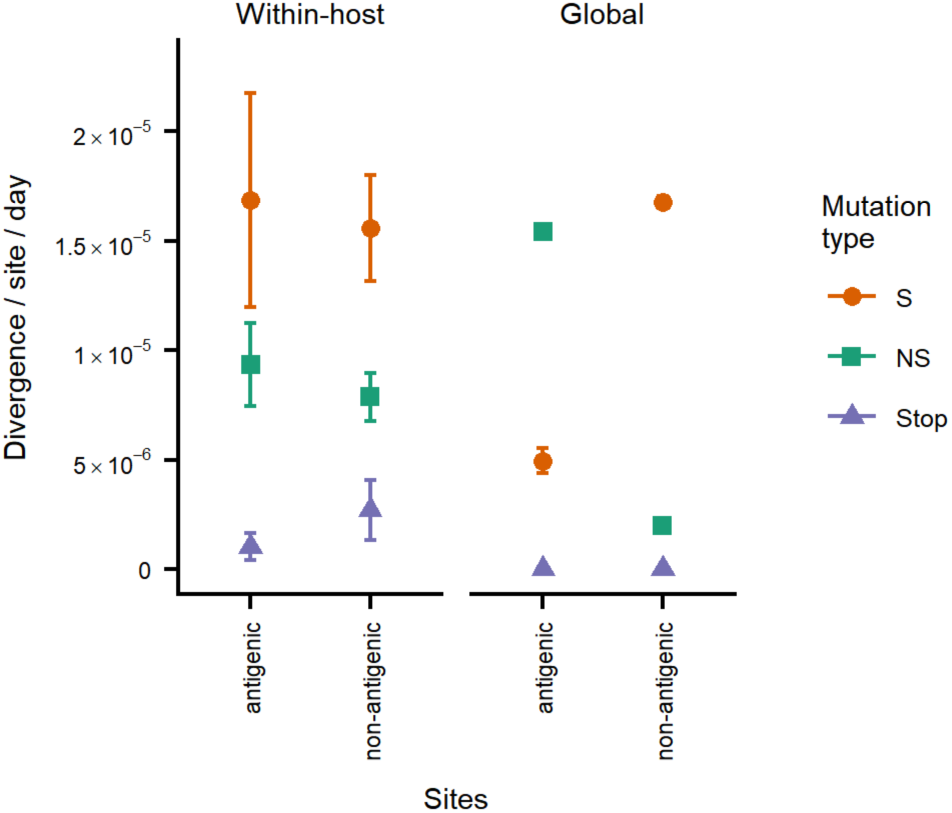
Antigenic and non-antigenic regions of hemagglutinin show similar patterns of variation within hosts, but antigenic regions evolve faster at the global scale. Within-host and global evolutionary rates were calculated as described in Figure 3 for a set of previously defined antigenic sites (Wolf et al., 2006). Within-host rates are shown as the mean and standard error of all patient infections sequenced in the three datasets analyzed after removing outlier samples (see **Materials and methods**)(Debbink et al., 2017; Dinis et al., 2016; McCrone et al., 2018). Global rates are shown as the mean and standard error calculated through linear regression (in most cases, the standard errors are smaller than the point sizes). Global rates of stop-codon evolution are zero because no stop codons are observed in patient consensus sequences.

**Figure 5.**
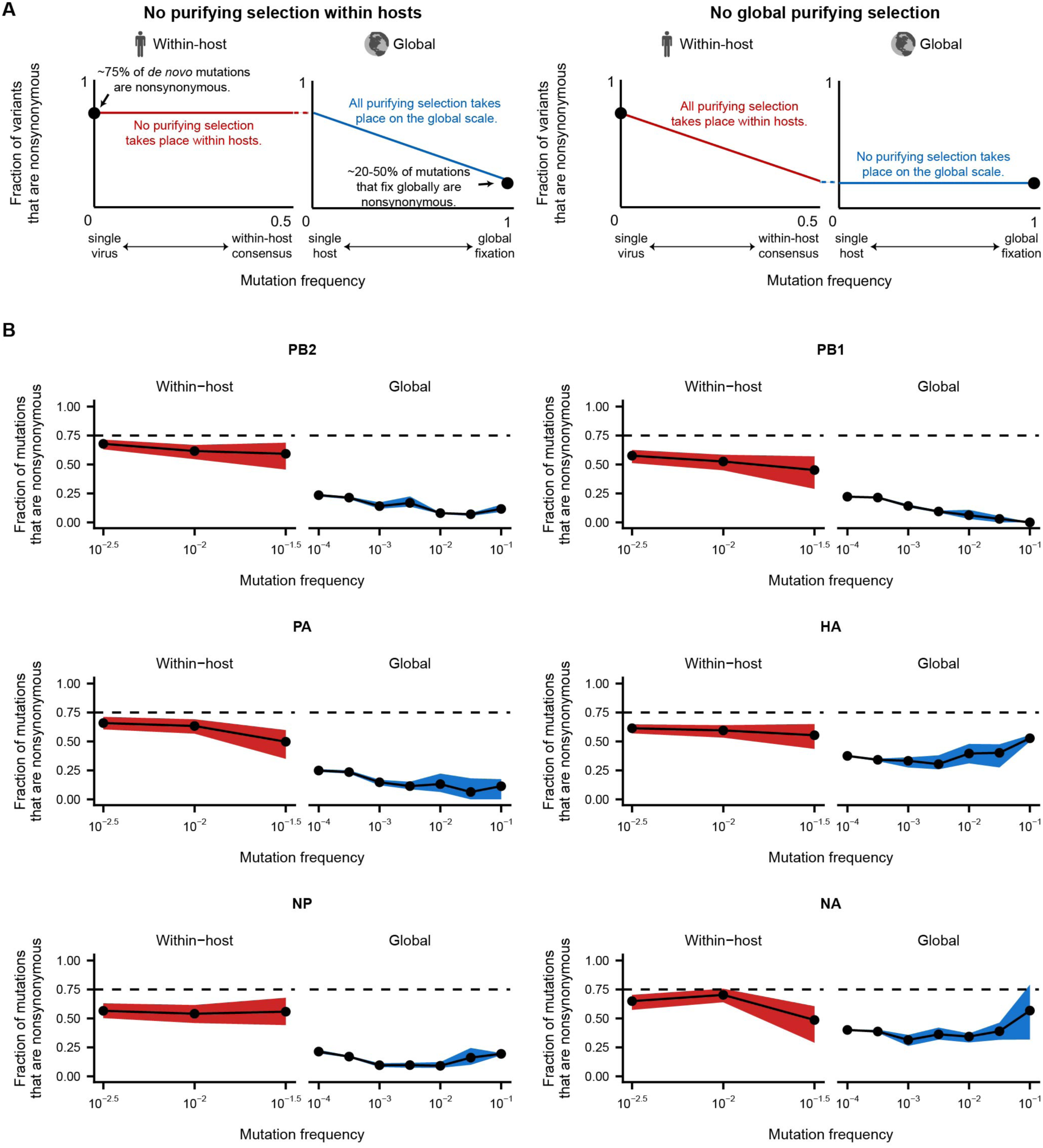
Within-host viral populations harbor many transient deleterious mutations that are purged from the global influenza population. **A)** The fraction of within-host and global mutations that are nonsynonymous reveals selective pressures that act across different evolutionary scales. In the absence of selection, about 75% of *de novo* mutations are expected to be nonsynonymous. However, only about 20-50% of the mutations that fix globally are nonsynonymous, suggesting that purifying selection acts within hosts and on globally circulating mutations to remove deleterious nonsynonymous variants. These plots illustrate two scenarios for how purifying selection acts on within-host and global viral populations. **B)** Fraction of within-host and global mutations that are nonsynonymous. Within-host viral populations harbor many nonsynonymous mutations, though less than would be expected in the absence of selection (dashed line). Purifying selection acts strongly at transmission and in the early stages of global evolution to lower the fraction of nonsynonymous mutations. Dashed lines indicate the ∼75% of *de novo* mutations that are expected to be nonsynonymous. The fraction of nonsynonymous mutations within hosts is shown as the mean and 95% confidence interval of 100 bootstrap samples of the within-host viral populations. The fraction of nonsynonymous mutations in the global influenza population is shown as the mean, minimum, and maximum of the values calculated separately for the 2015-2016, 2016-2017, and 2017-2018 H3N2 seasons.

To compare rates of evolution in acute and chronic influenza infections, we also calculated rates of influenza evolution in four chronic infections that lasted multiple months (Xue et al., 2017)(Figure S6). We find roughly similar rates of evolution in acute and chronic infections, except in hemagglutinin (HA) and neuraminidase (NA), which appear to evolve especially rapidly in these chronic infections due to selection for antigenic variation and antiviral resistance respectively. However, the small number of chronic infections constrains our interpretation.

### Purifying selection acts weakly within hosts to eliminate deleterious variants

What kinds of mutations reach detectable frequencies within hosts? We compared how quickly synonymous, nonsynonymous, and stop-codon (nonsense) mutations accumulate within hosts. Differences in how quickly these three classes of mutations accumulate can reveal how selection acts within hosts. Most synonymous mutations are nearly neutral; many nonsynonymous mutations are deleterious; and premature stop codons are lethal. In the absence of selection, all three types of mutations accumulate at the same rate. However, purifying selection purges deleterious mutations, and positive selection increases the frequency of any beneficial mutations.

We find that deleterious mutations are purged detectably but incompletely within infected individuals. Synonymous mutations accumulate about twice as quickly as nonsynonymous mutations within hosts, with some variation across genes (Figure 3C). Stop-codon mutations accumulate even more slowly within hosts than nonsynonymous mutations, but they are not completely purged. The depletion of nonsynonymous and stop-codon mutations relative to synonymous mutations demonstrates that viral populations within hosts experience purifying selection, as previous studies have observed (Dinis et al., 2016; McCrone et al., 2018; Moncla et al., 2019). However, the presence of stop-codon mutations at frequencies as high as 0.5% within hosts suggests that purifying selection remains incomplete, allowing some strongly deleterious mutations to persist long enough to reach detectable frequencies (McCrone et al., 2018). These results show that purifying selection acts detectably but weakly within hosts to reduce the frequencies of deleterious mutations.

### Synonymous mutations accumulate neutrally between hosts, but nonsynonymous mutations experience purifying and positive selection on a global scale

How do mutations that arise within hosts fare in the global viral population? We compared rates of evolution within hosts and globally to determine how selection acts across different scales of viral evolution. Neutral mutations should have similar rates of evolution within and between hosts. Purifying selection that acts between hosts to eliminate deleterious mutations will decrease global evolutionary rates compared to rates within hosts. Conversely, positive selection that acts on a global scale will increase rates of evolution between hosts relative to their within-host expectations for beneficial classes of mutations.

To calculate the global evolutionary rates of influenza virus, we analyzed H3N2 influenza sequences in the Global Initiative on Sharing All Influenza Data (GISAID) EpiFlu database, where each sequence represents the consensus sequence of one patient infection (Bogner et al., 2006). We used a molecular-clock method to estimate global evolutionary rates at synonymous and nonsynonymous sites in each gene (Figure 3B, Figure S7).

Synonymous mutations accumulate at similar rates within hosts and globally (Figure 3C). This concordance of evolutionary rates suggests that synonymous evolution in the global influenza population represents the simple accumulation of within-host variation through multiple patient infections, as expected for mutations that are nearly neutral (Kimura, 1968). Measurements of viral variation within hosts are noisy and incomplete due to technical limitations of deep sequencing (McCrone and Lauring, 2016), but this similarity in evolutionary rates between scales indicates that studies of within-host variation still capture representative slices of global variation.

In contrast, nonsynonymous mutations accumulate more slowly globally than within hosts in all six viral genes analyzed. In addition, while stop codon-mutations accumulate at appreciable rates within hosts, they never fix during global evolution. These discrepancies in evolutionary rates within and between hosts suggest that many nonsynonymous mutations that reach detectable frequencies within hosts are later purged by purifying selection as viruses circulate in the global influenza population.

An important exception to this trend occurs in the antigenic regions of the hemagglutinin (HA) gene, which evolve more rapidly on a global scale than non-antigenic regions of HA (Figure 4)(Caton et al., 1982; Fitch et al., 1997; Koel et al., 2013; Wolf et al., 2006). Within hosts, synonymous and nonsynonymous mutations accumulate at similar rates in HA antigenic regions as in the rest of the HA gene, suggesting that selection does not detectably favor antigenic mutations within hosts. However, nonsynonymous mutations in antigenic regions of HA accumulate more rapidly globally than they do within hosts. This observation suggests that antigenic selection at the between-host scale is responsible for amplifying the frequency of nonsynonymous antigenic mutations that arise within hosts.

### Transient deleterious variants within hosts are purged at transmission and in the early stages of global evolution

The previous section shows that nonsynonymous mutations are less common among mutations that fix globally than they are within hosts. This result suggests that purifying selection eliminates many nonsynonymous within-host variants before they fix in the global population of influenza viruses. However, it remains unclear from this analysis how quickly deleterious, nonsynonymous mutations that are generated within hosts are eliminated from the global viral population.

We tracked the fraction of nonsynonymous mutations at a range of mutation frequencies within and between hosts to determine how long deleterious, nonsynonymous mutations persist in the global viral population (Figure 5A). This analysis draws on the same logic that underlies the classical McDonald-Kreitman test for positive selection (Bhatt et al., 2010, 2011; Garud et al., 2019; McDonald and Kreitman, 1991; Rand and Kann, 1996). We expect purifying selection to purge deleterious nonsynonymous mutations from the viral population before they reach high mutation frequencies. In the absence of purifying selection, about 75% of *de novo* mutations are expected to be nonsynonymous. However, purifying selection reduces the fraction of nonsynonymous mutations by removing deleterious mutations after they arise and before they fix. By examining when nonsynonymous mutations are depleted from the global viral population, we can identify where in the evolutionary process most purifying selection takes place (Figure 5A).

Within hosts, about 55-70% of mutations were nonsynonymous in each influenza gene (Figure 5B). Because there are fewer nonsynonymous mutations within hosts than expected under *de novo* mutation alone, this result indicates that purifying selection acts detectably but weakly within hosts, supporting the conclusions of our analyses of evolutionary rates. Across a range of mutation frequencies within hosts, the fraction of nonsynonymous mutations remained relatively consistent, indicating that all mutations within hosts have experienced similar levels of purifying selection regardless of their mutation frequency.

In contrast, the fraction of nonsynonymous mutations drops sharply at transmission and in the early stages of global evolution. More than 50% of mutations within hosts are nonsynonymous, but only about 25% of low-frequency global mutations are nonsynonymous in the polymerase genes (PB2, PB1, PA, and NP). The fraction of nonsynonymous mutations is even lower among global mutations that reach higher frequencies, and eventually, fewer than 20% of the mutations that fix in these genes are nonsynonymous. Previous work has shown that rare variants in global viral populations are dominated by transient deleterious mutations (Pybus et al., 2007), so this decline in the proportion of nonsynonymous mutations at high frequencies likely occurs as purifying selection purges deleterious variation. Nevertheless, most deleterious nonsynonymous mutations are eliminated at transmission and the early stages of global evolution, before these mutations are detected by current global surveillance.

More nonsynonymous mutations are present globally in the viral surface proteins HA and NA, which experience strong positive selection for antigenic variation (Bhatt et al., 2011; Caton et al., 1982; Fitch et al., 1997; Koel et al., 2013; Monto et al., 2015; Murphy et al., 1972; Rambaut et al., 2008). In HA and NA, about 40% of rare global mutations are nonsynonymous. The fraction of nonsynonymous mutations increases slightly among more common mutations, and about 50% of mutations that eventually fix in the global population are nonsynonymous, as previously observed (Bhatt et al., 2011; Strelkowa and Lässig, 2012). We hypothesize that purifying selection drives the initial decline in nonsynonymous mutations at early stages of global evolution by purging deleterious mutations. Later, positive selection for antigenic variation likely increases the proportion of nonsynonymous mutations among commonly circulating strains. However, our analyses cannot distinguish the specific contributions of purifying and positive selection in shaping influenza’s global variation.

Our analyses show that transient deleterious mutations make up a large proportion of influenza’s genetic variation within hosts. Deleterious variation is common in global viral populations as well (Bhatt et al., 2011; Pybus et al., 2007) and can slow global rates of influenza virus evolution (Illingworth and Mustonen, 2012; Koelle and Rasmussen, 2015), but our findings show that deleterious mutations are substantially more prevalent within hosts. These results also show that purifying selection acts strongly to remove deleterious variation at transmission and in the early stages of global evolution as rare variation begins to circulate in the global influenza population.

## Discussion

In this study, we quantitatively compared influenza virus’s genetic diversity within and between human hosts. Our results show that although viral diversity may seem low within acute human influenza infections, it is sufficient to explain the virus’s rapid global evolution. Synonymous mutations accumulate at similar rates within and between hosts, as expected for neutral genetic changes. Most nonsynonymous mutations that reach detectable frequencies within hosts are eventually purged between hosts, suggesting that within-host influenza diversity consists mostly of transient, deleterious mutations that are eliminated at transmission and global evolution. Antigenic sites of HA are the only region of the influenza genome where mutations are consistently favored in the global viral population beyond their frequencies within hosts.

Our findings show how purifying and positive selection act on viral populations at different evolutionary scales. Influenza populations within hosts accumulate genetic variation as they expand during the first few days of an infection. However, most of this variation, which is deleterious, is purged as viruses circulate between hosts. One recent study estimates that influenza’s transmission bottleneck consists of one to two viral genomes (McCrone et al., 2018; Xue and Bloom, 2019), and further study is required to determine whether and how this narrow transmission bottleneck helps eliminate deleterious variants.

Our observations of a substantial excess of deleterious viral mutations within hosts extend prior work showing that globally circulating strains of influenza carry a deleterious mutational load that influences the dynamics of adaptation (Illingworth and Mustonen, 2012; Koelle and Rasmussen, 2015; Pybus et al., 2007; Strelkowa and Lässig, 2012). The nearly neutral theory of molecular evolution predicts that these transient, deleterious mutations will persist and sometimes even reach fixation when effective population sizes are small (Ohta, 1992). We demonstrate that the proportion of deleterious mutations is higher within hosts than it is globally, probably because within-host populations have small effective population sizes due to frequent transmission bottlenecks (McCrone et al., 2018). Co-infection and genetic complementation within hosts can also reduce the efficiency of selection by masking the effects of deleterious variation, as shown by the accumulation of defective interfering particles *in vitro* (Brooke, 2017; Davis et al., 1980; Frensing et al., 2013).

This finding of a high deleterious viral mutational load within human influenza infections agrees with analyses of other RNA viruses that cause acute infections. Studies of dengue (Holmes, 2003) and Lassa virus (Andersen et al., 2015) have likewise identified an excess of nonsynonymous mutations within hosts compared to global viral populations, suggesting that many RNA viruses with rapid mutation rates and frequent transmission bottlenecks may accumulate transient, deleterious variation within hosts.

Our work also clarifies how antigenic selection shapes influenza evolution within and between hosts. Antigenic mutations are not detectably enriched within hosts relative to other nonsynonymous mutations, suggesting that positive selection primarily acts between rather than within hosts to favor the mutations that drive influenza’s global antigenic evolution. Most antigenic selection may result from antibodies that prevent the initiation of new infections, and antigenic variants may be most strongly favored at transmission, where they are more likely to found new infections (Han et al., 2019; Petrova and Russell, 2018). However, positive selection within hosts that acts on antigenic mutations below our limit of detection may still increase the chance that antigenic variants transmit successfully and found new infections.

This finding that antigenic selection is limited in acute influenza virus infections contrasts with the rapid antigenic evolution that has been observed in chronic influenza virus infections, where multiple viral lineages carrying distinct antigenic mutations arise and compete within a single patient (McMinn et al., 1999; Xue et al., 2017). These differences in antigenic evolution between acute and chronic infections may reflect the fact that acute infections provide limited time for selection to have detectable effects on antigenic variation. Further study is required to identify the exact immune responses and mechanisms that drive influenza’s antigenic evolution.

Deep sequencing makes it possible to examine viral evolution at high resolution within natural human infections, but it has remained unclear how within-host genetic variation is transformed into the macroscopic evolutionary dynamics that occur at a global scale. Our work places influenza’s within-host genetic diversity in the context of its global evolution and provides a general framework for linking viral evolutionary dynamics across scales.

## Materials and methods

### Data and code availability

We downloaded raw sequencing data from the NCBI SRA database for Bioprojects PRJNA344659 (Debbink et al., 2017), PRJNA412631 (McCrone et al., 2018), and PRJNA364676 (Xue et al., 2017). We obtained sequencing data for (Dinis et al., 2016) by personal communication. The computer code that performs the analysis is available at https://github.com/ksxue/within-vs-between-hosts-influenza. Sequences downloaded from the GISAID Epiflu database are not available due to data-sharing restrictions, but acknowledgement tables for the sequences we analyzed are available at https://github.com/ksxue/within-vs-between-hosts-influenza/tree/master/data/GISAID/acknowledgements/H3N2.

### Variant calling and annotation

Here, we summarize our general variant-calling pipeline, with study-specific modifications described in more detail below. We used cutadapt version 1.8.3 to trim Nextera adapters, remove bases at the ends of reads with a Q-score below 24, and filter out reads whose remaining length was shorter than 20 bases (Martin, 2011). To determine the subtype of each sample, we used bowtie 2 version 2.2.3 on the --very-sensitive-local setting to map 1000 reads from each sample to reference genomes for each subtype: A/Victoria/361/2011 (H3N2), A/California/04/2009 (pdmH1N1), and A/Boston/12/2007 (seasonal H1N1)(Langmead and Salzberg, 2012). We classified the subtype of each sample based on which reference genome resulted in the highest mapping rate. For this study, we only analyzed samples of H3N2 influenza, which were the most common subtype sequenced in the datasets we analyzed.

For each sample, we mapped sequencing reads against the subtype reference genome using bowtie2 on the --very-sensitive-local setting (Langmead and Salzberg, 2012). We tallied the counts of each nucleotide at each genome position and inferred the sample consensus sequence using custom scripts. We then re-mapped all reads from each sample against the sample consensus sequence using the --very-sensitive-local setting of bowtie2, and we removed duplicate reads using picard version 1.43 from the GATK suite (McKenna et al., 2010). We tallied the counts of each nucleotide at each genome position using custom scripts, discarding reads with a mapping score below 20 and bases with a Q-score below 20. We annotated mutations as synonymous or nonsynonymous based on their effect in the background of the sample consensus sequence.

We defined sites of putative within-host variation as positions in the genome with sequencing coverage of at least 400 reads at which a minority base exceeded a frequency of 0.5%. In some cases, we also assessed the effects of using variant-frequency thresholds of 1% and 2% (Figure S2).

#### Study-specific modifications

We obtained sequencing data from the (Dinis et al., 2016) study in the form of a single FASTQ file per sample containing first and second members of read pairs. We reconstructed read pairs for each sample using read-pair information in the FASTQ headers, and then we analyzed the reconstructed pairs as described above.

### Sample exclusions

We first identified outlier samples with large amounts of within-host variation (Figure S3), since this high variation could be generated through co-infection or through poor sample quality and low viral load. For each study, we identified the top 10% of samples based on the number of within-host variants above a frequency threshold of 0.5%, and we excluded these samples from subsequent analyses. We also excluded samples that did not meet our minimum genome-coverage requirement that >80% of the sites in each sequenced gene have ≥400x coverage, as well as plasmid controls.

The (McCrone et al., 2018) study sequenced 43 longitudinal pairs of samples. Each pair was collected from the same subject during a single illness, and the samples in a pair were collected 0 to 6 days apart. The first sample in each pair was typically collected by the study subject in a home setting, and the second sample was collected in a clinical setting. We expect samples in a longitudinal pair to have correlated patterns of viral diversity, so we excluded the earlier sample in each longitudinal pair.

After excluding these samples, we analyzed the remaining 308 of the original 411 samples of acute H3N2 influenza in these three datasets.

### Within-host rates of evolution

We calculated the average divergence of each viral population from its consensus sequence by summing the frequencies of synonymous, nonsynonymous, and stop-codon (nonsense) mutations within hosts in each influenza gene (Figure 3A). Random mutations to the genome are more likely to be nonsynonymous than synonymous because of the structure of the genetic code, so we normalized sample divergence by the number of sites available for each type of mutation. We counted the proportion of sites available for synonymous, nonsynonymous, and nonsense mutations in the A/Victoria/361/2011 (H3N2) reference genome using a modified version of the Nei and Gojobori method that tallies nonsense mutations separately from nonsynonymous mutations (Nei and Gojobori, 1986). In this estimate, we assumed that transitions are about three times as common as transversions based on previous studies (Bloom, 2014; Pauly et al., 2017; Sanjuán et al., 2010). Using these methods, we calculated that 72% of mutations would be nonsynonymous, 25% synonymous, and 3% nonsense. To calculate the number of sites available for each type of mutation, we multiplied the length of the gene by the proportion of available sites. We then divided the viral divergence in each gene by the number of available sites to obtain a per-site viral divergence. We excluded the overlapping M1/M2 and NS1/NEP pairs of genes from all subsequent analyses because their out-of-frame overlap makes it possible for a single mutation to have multiple effects in the two genes.

To calculate rates of viral divergence per day, we normalized per-site viral divergence by the timing of sample collection. Of the 308 H3N2 samples that we analyzed, 251 had metadata describing the number of days post-symptom-onset after which the sample was collected. For the remaining samples sequenced by (Dinis et al., 2016), we made use of aggregate data on the timing of sample collection relative to symptom onset for all H3N2 samples in the study, as we did not have access to data on the timing of sample collection for individual samples. The average timing of sample collection varied across studies (Figure S1). Symptom onset typically occurs 2-3 days after infection begins (Baccam et al., 2006; Beauchemin and Handel, 2011; Carrat et al., 2008), so we added two days to the number of days post-symptom-onset to obtain the number of days post-infection (DPI). We calculated rates of viral divergence by dividing each sample’s per-site divergence by its DPI. We then calculated the mean and standard error of the viral divergence rates across all samples to obtain the values in Figure 3C.

For comparison, we calculated rates of evolution in acute patient infections through both the “point” method described above, which averages rates of evolution calculated from each patient infection, as well as through linear regression (Figure S5). We performed linear regression of per-site viral divergence by the sample DPI. Because we did not have access to sample-specific data on the timing of sample collection for the (Dinis et al., 2016) dataset, we excluded the (Dinis et al., 2016) samples from our estimate of evolutionary rates through linear regression.

To calculate within-host rates of evolution in antigenic sites of HA, we used the antigenic sites defined by (Wolf et al., 2006), and we tallied the synonymous, nonsynonymous, and nonsense mutations at those codons.

### Between-host evolutionary rates

We downloaded all full-length H3N2 influenza coding sequences in the Global Initiative on Sharing All Influenza Data (GISAID) EpiFlu database collected from January 1, 1999 to December 31, 2017 (Bogner et al., 2006). We randomly subsampled sequences for each gene to a maximum of 50 per year, we aligned these sequences to the coding regions of reference strains A/Moscow/10/1999 (H3N2) and A/Brisbane/10/2007 (H3N2) using the default settings of mafft version 7.407 (Katoh and Standley, 2013), and we trimmed non-coding regions from each sequence. We calculated the synonymous and nonsynonymous distance between each sequence and both the Moscow/1999 and Brisbane/2007 reference sequences using custom scripts. We excluded outlier sequences whose distance substantially exceeded the distances of other sequences in that year, since these outlier sequences may have been misannotated. As in our calculation of within-host rates of evolution, we excluded the overlapping M1/M2 and NS1/NEP pairs of genes because their out-of-frame overlap makes it possible for a single mutation to have multiple effects in the two genes.

We used a molecular-clock approach to calculate global evolutionary rates. We performed linear regression of the distances from a reference sequence by the timing of sample collection to estimate the rate of between-host evolution (Figure 3B, Figure S7). Molecular-clock methods can underestimate evolutionary rates over long periods of time as multiple mutations begin to occur at the same sites. To limit the effect of multiple-hit mutations on our analyses, we analyzed sequences collected over shorter intervals for more rapidly evolving genes. For the HA and NA genes, which evolve rapidly (Rambaut et al., 2008), we analyzed sequences from 2007 to 2017 relative to the Brisbane/2007 reference sequence. For the other genes, we analyzed a larger sequence set collected from 1999 to 2017 relative to the Moscow/1999 reference sequence. To calculate between-host rates of evolution in antigenic sites of HA, we used the antigenic sites defined by (Wolf et al., 2006), and we tallied the synonymous and nonsynonymous mutations at those coding sites.

### Chronic evolutionary rates

We previously deep-sequenced longitudinal samples of influenza from four chronic H3N2 influenza infections (Xue et al., 2017). We downloaded deep-sequencing data for these chronic infections from BioProject PRJNA364676. We mapped reads and identified within-host variants above a frequency of 0.5% as described above, except that we calculated within-host variant frequencies and annotated mutation effects relative to the consensus sequence of the influenza population at the first sequenced timepoint for each patient. After calling within-host variants, we calculated per-site viral divergence at each sequenced timepoint as described above, and we performed linear regression of per-site viral divergence by the time elapsed since the infection began for each patient. To obtain the aggregate evolutionary rates shown in **Figure S6A**, we calculated the mean and standard error of the evolutionary rates estimated for each patient.

### Fraction of mutations that are nonsynonymous within and between hosts

To calculate the fraction of mutations that are nonsynonymous within hosts, we tallied the total number of synonymous and nonsynonymous mutations in each frequency bin for each sample. Stop-codon mutations were counted as nonsynonymous mutations in this analysis. We then summed the variants in each frequency bin across samples. We calculated confidence intervals by performing 100 bootstrap resamplings of the viral samples and plotted the 95% confidence interval from these bootstrap replicates (Figure 5B).

To calculate the fraction of mutations that are nonsynonymous between hosts, we must be able to determine when the same mutation has arisen multiple times in the global influenza population. We downloaded all full-length H3N2 influenza coding sequences in the Global Initiative on Sharing All Influenza Data (GISAID) EpiFlu database collected from July 1, 2015 to June 30, 2018, together representing the 2015-2016, 2016-2017, and 2017-2018 Northern Hemisphere flu seasons (Bogner et al., 2006). We grouped all strains into seasons beginning on July 1 and ending on June 30 and analyzed each season separately. Passaged sequences can generate false signals of positive selection due to lab-adaptation mutations (McWhite et al., 2016), so we retained only unpassaged sequences for downstream analyses. We aligned sequences from each season and each gene to the coding regions of reference strain A/Victoria/261/2011 (H3N2) using the default settings of mafft version 7.407 (Katoh and Standley, 2013), and we trimmed non-coding regions from each sequence. We built a maximum-likelihood phylogeny for each gene and season using RAxML version 8.2.3 and the GTRCAT model (Stamatakis, 2014). We rooted the resulting phylogenies with A/Victoria/261/2011 as the outgroup, and we reconstructed ancestral states at each node using TreeTime (Sagulenko et al., 2018). We wrote custom scripts that used the Bio.Phylo Biopython package (Talevich et al., 2012) to traverse each tree, identify the mutations at each node, assess whether the mutations had synonymous or nonsynonymous effects, and count the number of descendants from that node. For each mutation at a node, we also identified cases in which a second mutation occurred at the same codon in the descendent sequences, and we subtracted the descendent sequences carrying the second mutation from the total number of descendants of the original node. We calculated the frequencies of between-host variants by dividing the number of descendants by the total number of sequences in that season. Shaded intervals in Figure 5B display the range of the proportion of nonsynonymous variants in the three seasons analyzed.

## Acknowledgements

We thank Tom Friedrich, Jorge Dinis, Kelsey Florek, Katarina Braun, Ed Belongia, and Jenny King for sharing the sequencing data and sample metadata for their earlier study, as well as Louise Moncla and Trevor Bedford for their helpful suggestions regarding these analyses. We also thank Allie Greaney, Louise Moncla, and Seungsoo Kim for their comments on the manuscript. K.S.X. was supported by the Hertz Foundation Myhrvold Family Fellowship. J.D.B. was supported by grant R01AI127893 from the NIAID and as an Investigator of the Howard Hughes Medical Institute.

## Supplemental figures

**Figure S1.**
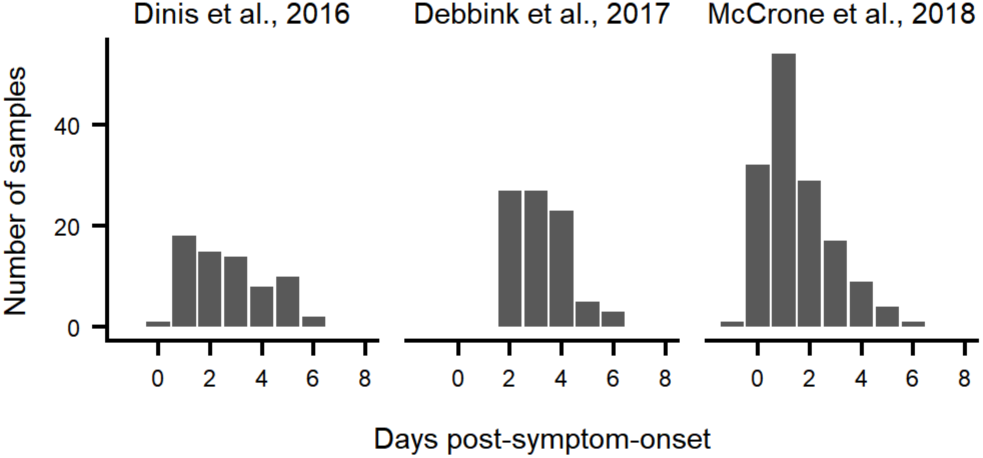
Distribution of days post-symptom-onset on which viral samples were collected. Note that symptoms typically emerge about 2 days after viral infection begins (Baccam et al., 2006; Carrat et al., 2008). Samples from the (Debbink et al., 2017) and (McCrone et al., 2018) studies that were excluded from analysis (see **Materials and methods**) were omitted from these distributions. All samples from the (Dinis et al., 2016) study are shown here, even though some of these samples were excluded from subsequent analyses, because metadata on the timing of sample collection was only available for all H3N2 samples in aggregate rather than for each sample individually.

**Figure S2.**
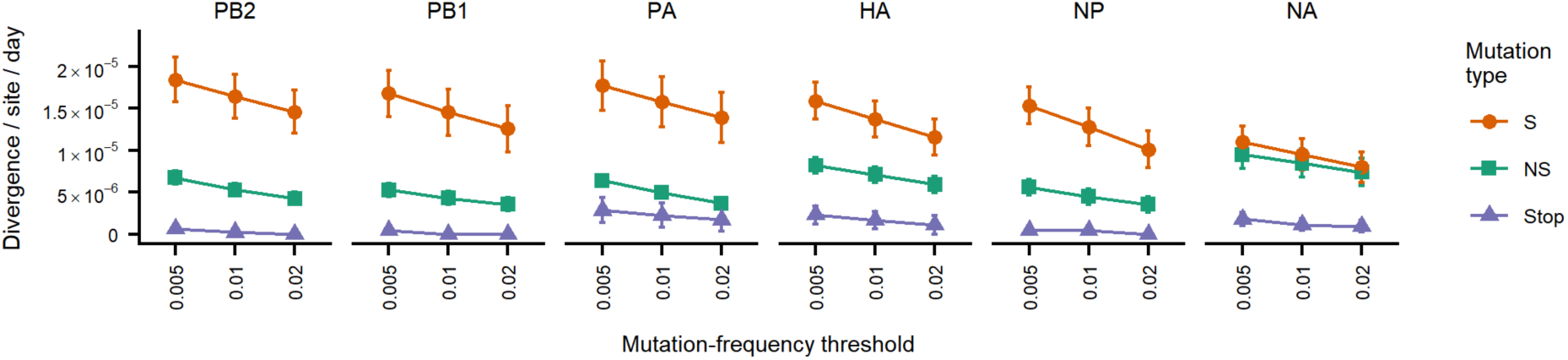
Estimates of within-host evolutionary rates are robust to variant-frequency thresholds. Shown are the mean and standard error of within-host evolutionary rates calculated as described in Figure 3 for commonly used variant-frequency thresholds.

**Figure S3.**
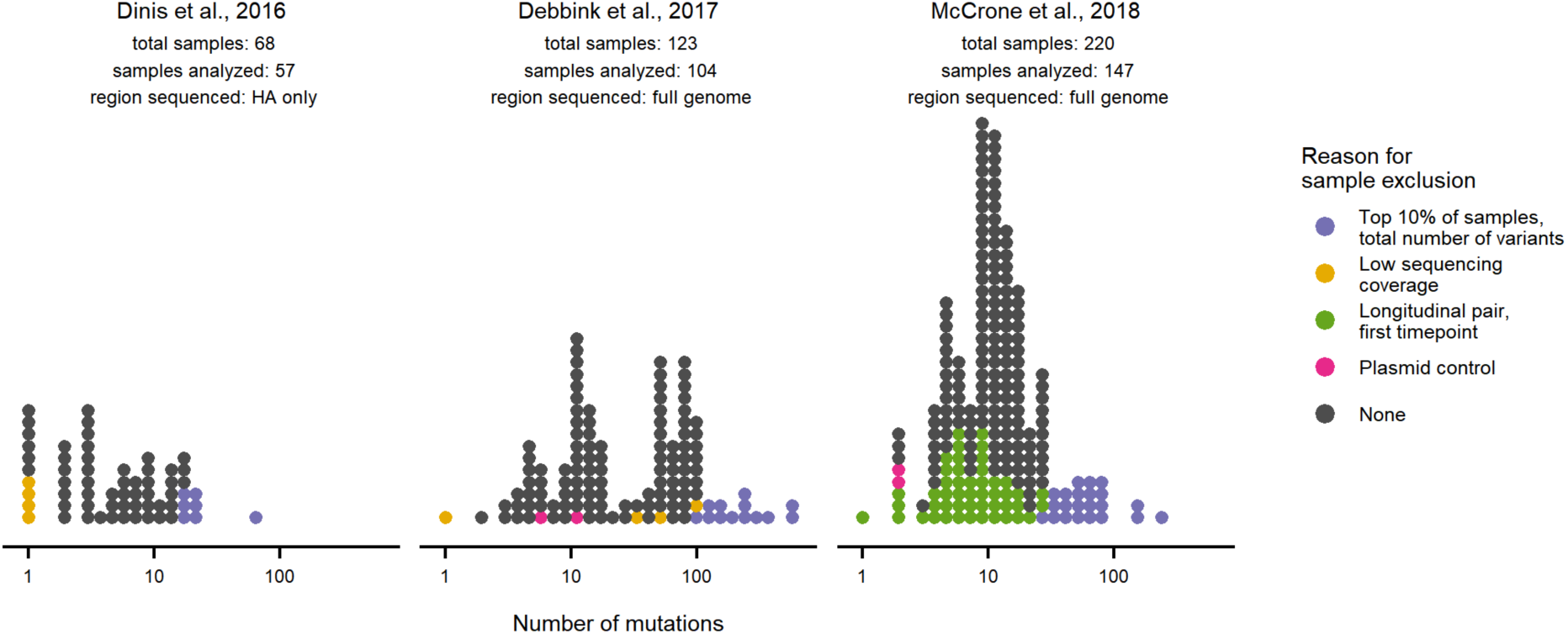
Samples excluded from downstream analyses. Each point represents the number of mutations identified above a frequency of 0.5% in a patient sample. Samples colored in grey were included in subsequent analyses. Samples in purple ranked in the top 10% of samples sequenced in that study based on the number of within-host mutations. Samples in yellow had incomplete sequencing coverage, meaning that fewer than 80% of sites in at least one gene were sequenced to 400x coverage. Samples in green were the first members of longitudinal pairs of samples obtained from the same patient infection; the first sample in each longitudinal pair was excluded to remove potential correlations in viral diversity between samples from a single patient. Samples in pink were plasmid controls that do not represent clinical viral populations.

**Figure S4.**
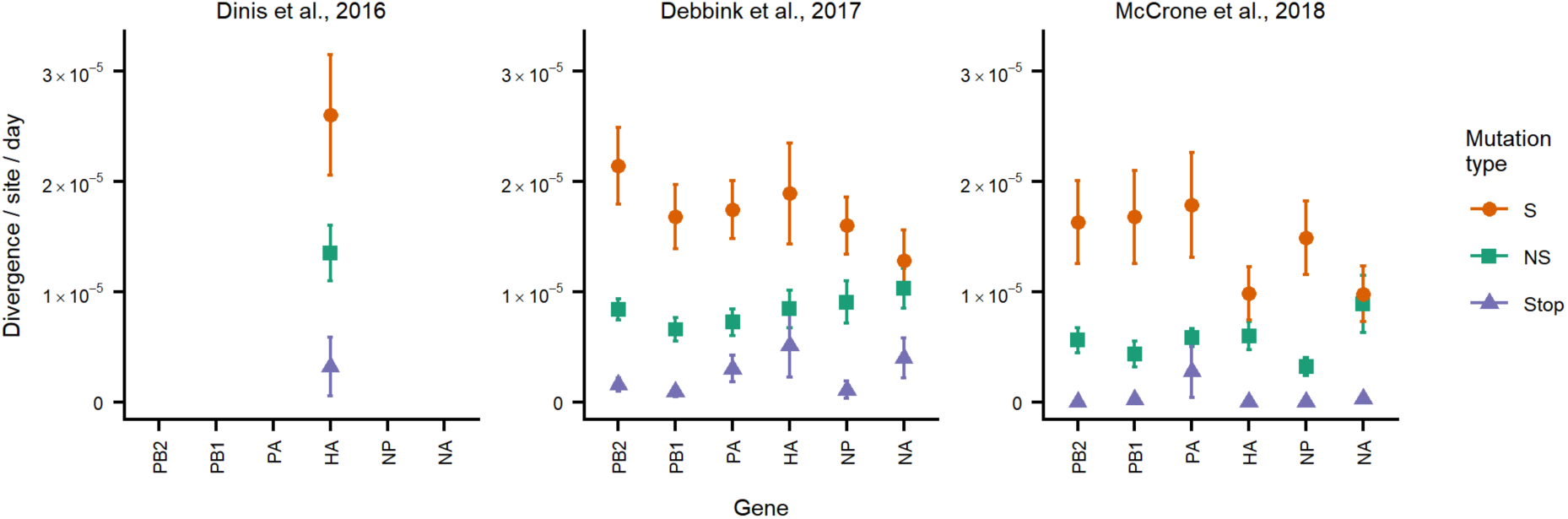
Estimates of within-host evolutionary rates are broadly consistent across cohorts. Shown are the mean and standard error of within-host evolutionary rates calculated as described in Figure 3 for viral samples in each published dataset. Note that (Dinis et al., 2016) sequenced only the HA gene.

**Figure S5.**
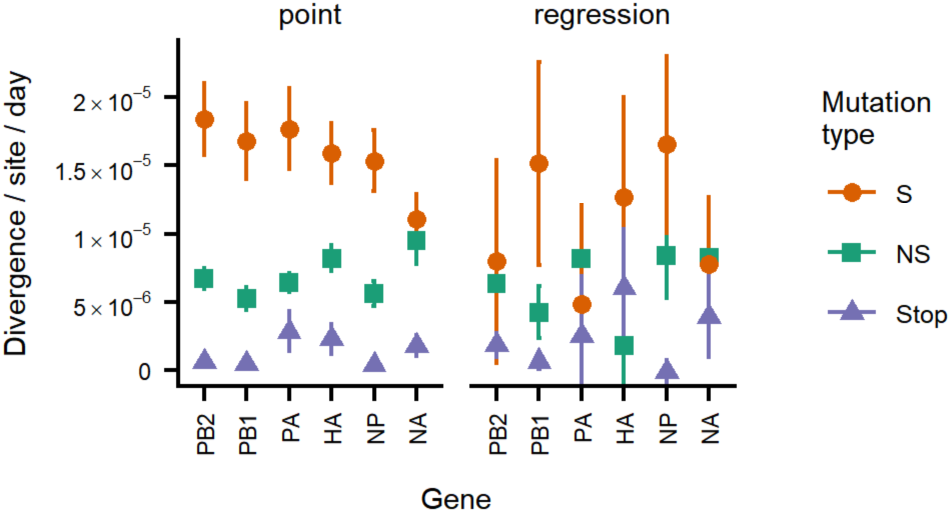
Within-host evolutionary rates show qualitatively similar trends when calculated using different methods. Evolutionary rates estimated using the point method are described and shown in Figure 3. Evolutionary rates were estimated using the regression method by calculating the total divergence of each within-host viral population, normalizing to the number of available sites, and then performing linear regression of per-site viral divergence by the time elapsed since each infection began. Rates estimated using the regression method do not include samples from the (Dinis et al., 2016) study because metadata on the timing of sample collection was only available for samples in aggregate for this dataset rather than the samples individually. (Rates estimated using the point method use aggregate metadata on the timing of sample collection.)

**Figure S6.**
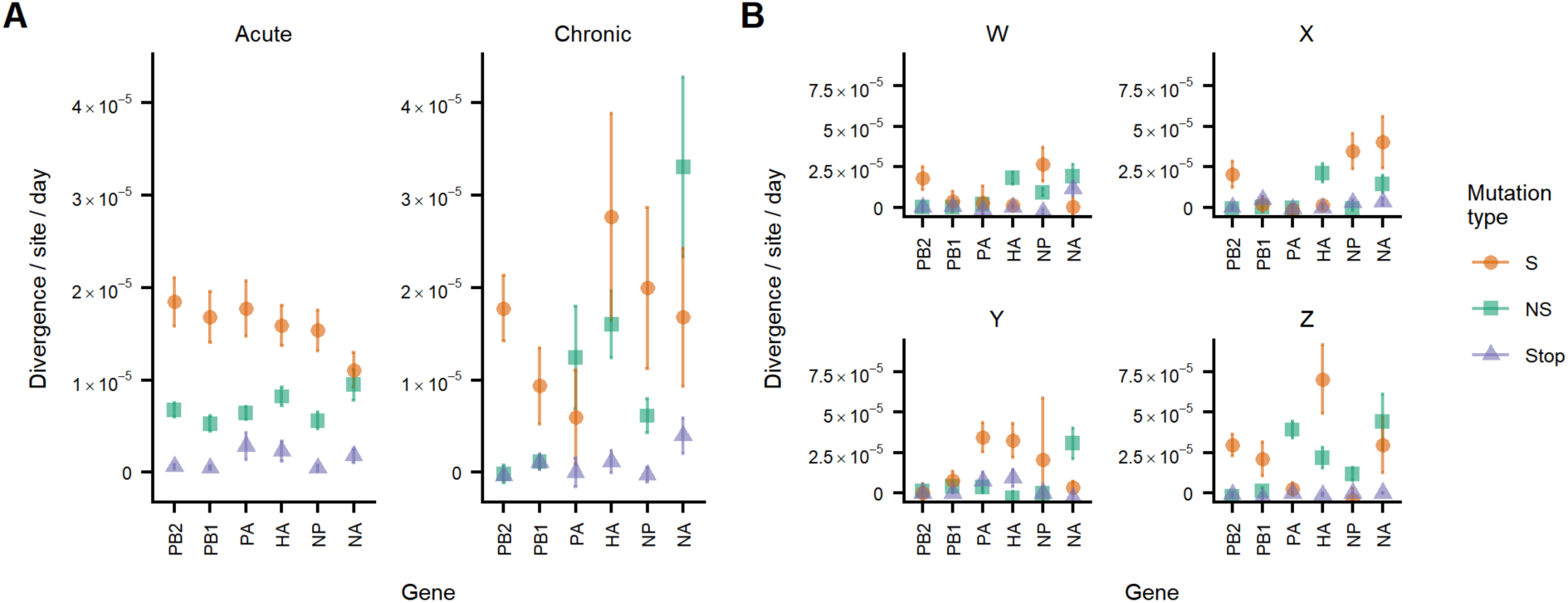
Within-host evolutionary rates in chronic influenza infections. **A)** Within-host evolutionary rates in acute and chronic infections. Evolutionary rates in acute infections were calculated as in Figure 3. Evolutionary rates in chronic infections were estimated separately for each of four patients from previously sequenced longitudinal viral samples (Xue et al., 2017) by calculating the total divergence of viral populations at each time point, normalizing to the number of available sites, and performing linear regression of per-site viral divergence by time since the infection began (see **Materials and methods**). Shown here are the mean and standard error of the evolutionary rates estimated for each patient. **B)** Within-host evolutionary rates plotted separately for each patient. Shown are the mean and standard error of evolutionary rates estimated as described above through linear regression. Patients are named as in (Xue et al., 2017).

**Figure S7.**
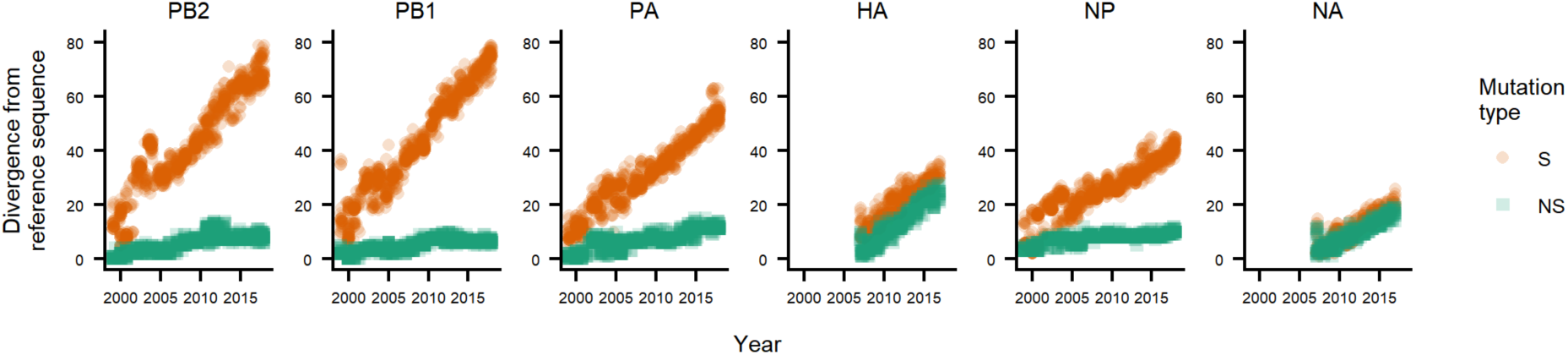
Estimate of global evolutionary rates using a molecular-clock method. Synonymous and nonsynonymous sequence divergences from a reference sequence are shown for randomly sampled sequences from the GISAID database (Bogner et al., 2006). For the PB2, PB1, PA, and NP genes, sequences from 1999-2017 were analyzed relative to a A/Moscow/10/1999 reference sequence. For the HA and NA genes, which evolve rapidly and can quickly saturate available sites of mutation, sequences from 2007-2017 were analyzed relative to a A/Brisbane/10/2007 reference sequence. Outlier sequences, which likely result from mis-annotations, were removed prior to performing this analysis as described in **Materials and methods.**

